# Microglial Purinergic Signaling Underlies Salt-Induced Neurovascular Polarity Reversal in the Hypothalamus During Heart Failure

**DOI:** 10.1101/2025.09.11.675347

**Authors:** Ranjan K. Roy, Elba Campos Lira, Manuel Bita Ongolo, Jessica A. Filosa, Javier E. Stern

## Abstract

**Background:** Neurovascular coupling (NVC) is essential for matching cerebral blood flow (CBF) to neuronal activity. While cortical NVC has been studied extensively, particularly in the context of sensory processing, little is known about NVC dynamics in deep brain regions, such as the hypothalamus, especially under disease conditions like heart failure (HF), where impaired cortical NVC has been linked to cognitive decline. Our goal in this study was to investigate salt-induced NVC responses in the hypothalamic supraoptic nucleus (SON) of rats with HF, and to determine the role of microglial purinergic signaling in modulating these responses.

**Methods:** Using *in vivo* two-photon imaging and real-time oxygen measurements in the SON, we assessed neurovascular responses to a systemic salt challenge in a well-established HF rat model that mimics clinical outcomes observed in the human population. Pharmacological and biosensor approaches were employed to dissect the contribution of key vasoactive mediators.

**Results:** Contrary to our original hypothesis, that HF would exacerbate salt-evoked inverse NVC (iNVC; vasoconstriction and hypoxia) as previously reported by our group in healthy rats, in HF, the NVC response was reversed. Here, salt-induced neuronal activation triggered vasodilation and increased SON pO₂, restoring oxygen levels to those of sham controls. This vasodilation was mediated by adenosine acting on A2A receptors and originated from a putative microglial source. Importantly, a masked, enhanced AVP-mediated vasoconstrictive component was still present, as revealed by biosensor assays, indicating a complex interplay between opposing vasoactive signals during HF.

**Conclusions:** These findings reveal a previously unrecognized microglia-driven purinergic mechanism that overrides AVP-mediated vasoconstriction to restore SON oxygenation during salt challenges in HF. The polarity switch in hypothalamic NVC suggests a region- and disease-specific adaptation with potential relevance to neurohumoral dysregulation in HF.

## INTRODUCTION

Neurovascular coupling (NVC) is a well-established phenomenon in the brain that involves a tight coupling between neuronal activity and increased cerebral blood flow (CBF) ^1,2^. This activity-dependent increase in CBF (also known as functional hyperemia) occurs rapidly (<2 s) and in a spatially restricted manner ^3,4^, ensuring optimal delivery of glucose and oxygen to brain areas with increased metabolic demand. NVC is generally thought to be a uniform process across brain regions and is typically mediated by byproducts of synaptic activity ^5^. However, we recently demonstrated a unique form of NVC in the hypothalamic supraoptic nucleus (SON), in which increased neuronal activity of neurosecretory vasopressin (AVP) neurons, triggered by a systemic salt challenge, induced an activity-dependent vasoconstriction (termed inverse NVC or iNVC) that led to local tissue hypoxia and further increase in AVP neuronal excitability ^6^. Furthermore, we demonstrated that the iNVC was mediated by local somatodendritic release of AVP, and that this positive feedback loop played an important role in the ability of these neurons to homeostatically respond to the peripheral interoceptive challenge ^6^.

A compromised NVC function is typically observed in several neurodegenerative disorders, such as Alzheimer’s disease and other forms of dementia ^7^, contributing to cognitive deficits in these diseases. Notably, NVC is also compromised in CVDs, including heart failure (HF) ^8,9^, hypertension ^10,11^ and stroke ^12^. A recent study in a mouse model of HF showed an impaired NVC response in the somatosensory cortex in response to whisker stimulation that contributed to cognitive impairment and dementia ^13^. Moreover, a compromised NVC in the cerebral cortex was demonstrated in HF patients with both reduced ejection fraction (EF) and preserved EF ^8,14^. Still, whether NVC responses in other brain regions relevant to the pathology of HF also occur in this disease, such as the hypothalamic iNVC, remains unknown.

A signature pathophysiological process in HF is neurohumoral activation, which involves increased sympathoexcitatory output to the heart, kidneys, and the vasculature, along with the systemic release of neurohormones, including AVP. Importantly, neurohumoral activation is directly associated with overall morbidity, mortality, and survival rates in HF ^15^. Several hypothalamic mechanisms have been shown to contribute to increased neurohumoral drive in HF, including increased AVP intrinsic neuronal excitability ^16,17^, changes in the excitatory/inhibitory balance ^18^, increased expression and release of AVP ^16,19^, and blunted nitric oxide (NO) availability and actions ^17,20,21^. Still, whether NVC responses in the hypothalamus are altered in HF, thereby contributing to the exaggerated neurohormonal outflow in this condition, remains unexplored. Finally, dietary salt is a key factor contributing to volume overload and congestion in HF ^22,23^, and excessive dietary salt intake has been implicated in additional neurohumoral activation in both HF and hypertension ^22,24,25^. Thus, evaluating whether NVC responses to salt are altered in HF could shed light on alternative mechanisms by which salt could further contribute to complications in this prevalent cardiovascular disease. In this study, we used a combination of *in vivo* and *ex vivo* approaches to test the hypothesis that an imbalance in vasoconstrictive/vasodilatory signals in the hypothalamic SON leads to exacerbated salt-evoked iNVC responses and consequently hypoxia in rats with HF.

## MATERIALS AND METHODS

### Animals

Male and female heterozygous transgenic eGFP-AVP Wistar rats (250-400gm) were used for all experiments ^26^. Rats were housed in cages (2 per cage) under constant temperature (22 ± 2°C) and humidity (55 ± 5%) on a 12-h light cycle (lights on: 08:00-20:00). All performed experiments were approved by the Georgia State University and the Augusta University Institutional Animal Care and Use Committee (IACUC) and carried out in agreement with the IACUC guidelines. At all times, animals had *ad libitum* access to food and water, and every effort was made to minimize suffering and the number of animals used in this study.

### Heart failure surgery and echocardiography

To induce heart failure in rats, a coronary artery ligation surgery was performed as previously described ^27^. Anesthesia was induced using 5% isoflurane mixed with O2 (100% O2, 1L/min). Optimum anesthesia was assessed by the absence of limb withdrawal reflex to a painful stimulus (hind paw pinch). Rats were then intubated for mechanical ventilation. During the surgery, anesthesia was adequately maintained using 2-3% isoflurane delivered by a vaporizer machine mixed with O_2_ (100% O_2_, 1L/min). With aseptic precaution, a small incision was made in the third intercostal space, and a retractor was applied to retract the 3rd and 4th ribs. The pericardium was ruptured, and the heart was exteriorized. A ‘Prolene 6-0’ suture (Ethicone, USA) was inserted just beneath the left atrial base along the interventricular septum to make a loop around the main diagonal branch of the left anterior descending (LAD) coronary artery. Finally, myocardial infarction was induced by ligating the LAD coronary artery. Occlusion of the LAD coronary artery results in a pale discoloration of the left ventricle, which guides as a visual confirmation of the ischemic left ventricle following LAD coronary artery ligation. Sham animals underwent a similar aseptic surgical procedure except for left coronary artery ligation. Four to five weeks after the surgery, we performed transthoracic echocardiography (Vevo 3100 systems; Visual Sonics, Toronto, ON; Canada) under light isoflurane (2-3%) anesthesia to assess the ejection fraction (EF) and confirm the development of HF. We obtained the left ventricle internal diameter and the left diameter of the ventricle posterior and anterior walls in the short-axis motion imaging mode to calculate the EF. The myocardial infarct surgery typically results in a wide range of functional HF, as determined by the EF measurements. Rats with EF<40% were considered as HF (Sham 81.68 ± 2.75%, HF 28.54 ± 4.58; n= 20 and 25 in sham and HF rats, respectively, see Table 1 for additional echocardiographic parameters). Rats that underwent HF surgery but did not develop HF or displayed an EF>40% were not included in the study.

**Table 1:** Summary of assessed cardiac parameters in sham and heart failure (HF) rats via echocardiography.

|  | **Sham**<br>**(n=20)** | **HF**<br>**(n= 25)** | **P value**<br>**(unpaired t-test)** |
| --- | --- | --- | --- |
| **Heart rate (BPM)** | 296.94 ± 27.22 | 286.85 ± 19.82 | P = 0.195 |
| **Stroke Volume (uL)** | 231.66 ± 25.79 | 137.28 ± 8.59 | P < 0.001 |
| **Ejection Fraction (%)** | 81.68 ± 2.75 | 28.54 ± 4.58 | P < 0.001 |
| **Fractional Shortening (%)** | 48.25 ± 2.31 | 15.48 ± 1.26 | P < 0.001 |
| **Cardiac Output (mL/min)** | 69.13 ± 6.59 | 39.02 ± 3.61 | P < 0.001 |
| **LV Mass (mg)** | 1832.49 ± 71.69 | 995 ± 51.29 | P < 0.001 |
| **Lvaw,d (mm)** | 3.16 ± 0.24 | 1.38 ± 0.15 | P < 0.001 |
| **Lvaw,s (mm)** | 3.82 ± 0.16 | 1.45 ± 0.16 | P < 0.001 |
| **Lvid,d (mm)** | 6.87 ± 0.24 | 12.38 ± 0.69 | P < 0.001 |
| **Lvid,s (mm)** | 4.81 ± 0.38 | 8.14 ± 0.36 | P < 0.001 |
| **Lvpw,d (mm)** | 4.10 ± 0.41 | 1.45 ± 0.25 | P < 0.001 |
| **Lvpw,s (mm)** | 3.97 ± 0.16 | 1.96 ± 0.16 | P < 0.001 |

Abbreviations: left ventricular end-diastolic anterior wall thickness (LVAW d), left ventricular end-systolic anterior wall thickness (LVAW s), left ventricular end-diastolic posterior wall thickness (LVPW d), left ventricular end-systolic posterior wall thickness (LVPW s), left ventricular end-diastolic diameter (LVID d), left ventricular end-systolic diameter (LVID s). All values in mm

### Surgery for *in vivo* 2-photon imaging of the rat supraoptic nucleus (SON)

A modified transpharyngeal surgical approach was used to expose the ventral surface of the brain containing the hypothalamic SON ^6,28^. Briefly, rats were anesthetized by intraperitoneal injection of urethane (1.5 gm/Kg) (U2500, Sigma Aldrich, MO, USA). Upon cessation of the hindlimb withdrawal reflex, the trachea and left femoral vein were catheterized, and the rats were placed supine on a heating pad in a stereotaxic frame (Stoelting 51600 Lab Standard Stereotaxic Instrument). After the oral cavity was opened by splitting the mandibles, cautery (Thermal Cautery Unit®, Geiger Medical Technologies, IA, USA) and drilling were used to remove the molars, ventromedial aspect of the right temporal bone, the basisphenoid and presphenoid bones to expose the ventral surface of the hypothalamus and ventral aspect of the right temporal lobe. The meninges overlying the right SON were removed, and a stainless-steel ring (i.d. 3.6 mm) was placed on the surface of the brain with the junction of the internal carotid artery (ICA) and middle cerebral artery (MCA) at the center.

All the drugs used were topically delivered through an MD probe (infusion rate 0.6ml/hour), which only penetrates a short distance inside the brain surface ^29^. To this end, a U-shaped microdialysis (MD) probe (permeable to 10 kDa; in-house design) ^30,31^ was bent horizontally at an angle of 100–150° and placed on the exposed surface of the SON, where the PA, ICA, and MCA were clearly visible inside the loop of the MD probe. In this process, drug concentrations achieved 0.5-1 mm below the brain surface are approximately four orders of magnitude lower compared to the concentration in the MD probe ^28^. Thus, to achieve a nanomolar range of drug concentration within the SON, drug concentration in the MD probe needs to be in the 1-2 mM range ^28^. The MD probe drug concentrations are noted in the corresponding results section.

### In-vivo 2-photon imaging of the SON and surrounding vasculature

The microvasculature of the SON and the surrounding area was labelled by i.v. injection of Rhodamine 70 kDa (20 mg/ml, 200 nl/rat) (R9379, Sigma Aldrich, MO, USA). The exposed ventral surface of the brain was covered with optical gel, and the SON and its surrounding vasculature were imaged under a 2-photon microscope (Bruker, Billerica, MA) excited with a Ti: Sapphire laser (Chameleon Ultra II [Coherent, Santa Clara, CA]) tuned at 840 or 940 nm and scanned with resonant galvanometers. As previously characterized ^6^, the visually accessible area of the SON is typically located medially to the bifurcation of the ICA and MCA. The SON was readily identified by the presence of eGFP-AVP-expressing neurons located within the dense capillary network. A schematic representation of the visually accessible area of the SON containing eGFP-AVP neurons and surrounding vascular cytoarchitecture is shown in ***Fig.1***. For measurements of vessel diameter and blood flow, we used either a 4X (numerical aperture 0.13) or a 16X (numerical aperture 0.8) water immersion objectives (Olympus, Center Valley, PA, USA), respectively.

**Figure 1.**
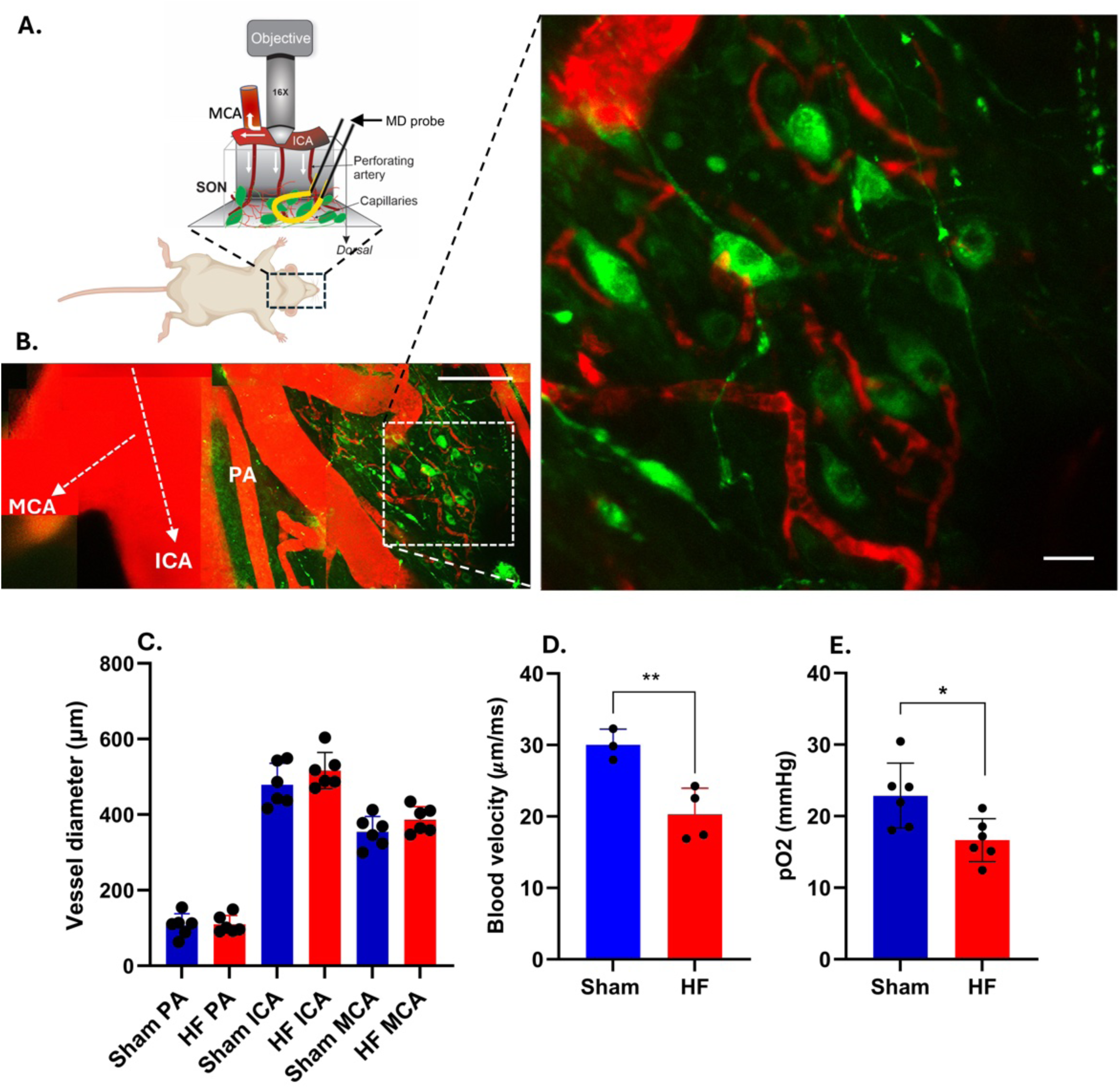
A persistent hypoxic milieu in the SON of HF rats. (**A**) Schematic representation of the preparation used for *in vivo* two-photon imaging of the SON. Note the presence of the microdialysis probe positioned right on top of the SON, used to locally deliver drugs into the region (**B**). Low-magnification (16x) two-photon image of the visually accessible area of the SON depicting main vessels in red following rhodamine 70 kDa administration (IV). The SON area is identified by the presence of a dense capillary network (red) along with eGFP-AVP-positive neurons (green), medially to the bifurcation of the internal carotid artery (ICA) and the middle cerebral artery (MCA). The magnified image to the right shows the capillary network and AVP cells from the area bounded by the white rectangle in the low magnification image to the left. (**C**) Bar graph summarizing the mean basal diameter of the parenchymal arteriole (PA), internal carotid artery (ICA), and the middle cerebral artery (MCA) in sham and HF rats (n=6 of each vessels/6 rats in each group). (**D**) Bar graph summarizing the mean blood velocity in SON PA of sham and HF rats. (n=3 in sham and 4 in HF group, **p<0.01; unpaired t-test). (**E**) Bar graph representing the mean oxygen saturation level (pO_2_) in the SON of sham and HF rats. (n=6 in each group, *p<0.05; unpaired t-test). Scale bar=100mm for low magnification and 25mm for high magnification. Error bars represent SEM.

### Quantification of vessel diameter and blood flow

Salt loading was performed via the femoral vein catheter by graded infusion of hypertonic saline (2M NaCl, 1 ml over 30 min) in HF and control (sham) rats. *<u>Vessel diameter</u>*: To quantify changes in vessel diameter during the salt loading challenge, a time-series, Z-stack acquisition protocol was used (1 iteration: 20 images at 0.33 Hz, 30 µm interval up/down the Z-axis in the SON; a total of 20 iterations with a 120 s interval; 512 x 512 pixels). Analyses were performed with *ImageJ,* as previously described^6^. Briefly, a Z-projected image was obtained for each of the 20 iterations, and vessel cross-section lines were manually drawn perpendicular to the two sides of the vessel wall. A profile of image brightness along the cross-section line of the vessel was then obtained. The high contrast between the vessel wall and the brain parenchyma allows for the ready detection of the vessel wall edges, which appear as a sudden rise in basal fluorescence. The distance between the two fast-rising phases, measured just above the base of the fluorescence profile, was used to calculate the diameter in each frame of the time series. Plots of these measurements over the time series were obtained to study the time course of diameter changes during the salt challenge. *<u>Blood flow</u>*: Blood flow was calculated by monitoring the movement of red blood cells (RBC) within the vessels, which appeared as moving dark spots in fluorescently labeled vessels through a segment of an imaged vessel. To this end, a separate set of experiments were performed, in which a line scan was performed at high speed (500-1000 Hz, 5-10 s) in a desired section of the targeted vessel. The line scans were readily converted into a kymographs by Prairie View software. In these graphs, the movement of RBCs appears as streaks in space-time images, in which the streak angle is a function of the RBC velocity ^6,32^. The kymograph generated from the line scan was then uploaded into Fiji software for the analysis of blood velocity ^33^. A line was manually drawn using the line tool along the streak angle to generate the angle of the line (the angle between the drawn line and the horizontal axis), taking care to consistently draw the line along the streak angle. This angle was then used to calculate the angle directly opposite the x-axis (i.e, distance). After converting this angle to radians, now represented as Ill, the following equation was used to calculate velocity:

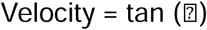

Once the velocity ratio was calculated, it was converted into μm/ms using the embedded distance (μm/pixel) and line period (ms/pixel) values from the line scan. This same procedure was repeated 20 times for each kymograph, sampling a different band each time to account for RBC velocity variability within the sampled time. The velocity values were then averaged for every line scan at each time point.

### Measurement of tissue partial pressure of oxygen (pO_2_)

The partial pressure of tissue O_2_ (pO_2_) in the SON was measured using an optical fluorescence probe (250 μm tip diameter; OxyLite system; Oxford Optronix), which allows for real-time recording of absolute changes in tissue pO_2_ as previously described ^6^. Rats were anesthetized with urethane (1.5 g/kg) and positioned in a stereotaxic apparatus. The SON was then exposed using a transpharyngeal surgery as described above. For these experiments, and to obtain a stable baseline pO_2_, rats were mechanically ventilated using a positive pressure ventilator (Harvard apparatus, Model-683, Holliston, MA, USA) with a tidal volume of ∼1 mL per 100 g of body weight and a frequency of 80 strokes/ min. The sensor was placed 1–2 mm below the ventral surface of the brain, medially to the bifurcation of the ICA and the MCA. The sensor was then retracted ∼0.1 mm to its final position to minimize tissue compression and allowed to equilibrate for at least 15–20 minutes before any measurements were obtained. SON pO_2_ was recorded at baseline at a rate of 1 Hz.

### Measurement of plasma osmolality

Blood was collected from the femoral vein after the salt challenge (2M NaCl, 1 ml i.v. over 30 min) or an isotonic saline infusion in sham and HF rats. Plasma was separated via centrifugation (2200-2500 RPM). The osmolality of the freshly obtained plasma was measured using a micro osmometer (Advanced Instruments, Model 3300, Version 4.3, Massachusetts, USA).

### Quantitative measurement of local AVP release within the SON using biosensor Sniffer^AVP^ cells

The full description of this methodological approach has been previously described ^34^. Briefly, Sniffer^AVP^ biosensors were generated by culturing Chinese hamster ovary cells in Dulbecco’s Modified Eagle Medium containing 10% w/v fetal bovine serum, 1% w/vpenicillin–streptomycin, 1% w/v Na-pyruvate and 1%w/v NaCO 3 filtered once through a Nalgene filtration system, and transfected with pcDNA3.1+ containing human V1a vasopressin receptors cloned in EcoRI (5’) and XhoI (3’) (plasmid obtained from Missouri S&T cDNA Resource Centre, Rolla, MO, USA) using lipofectamine, and stable over expression was achieved by geneticin (500 mg/ml) selection. About 16−20 h before the experiment, Sniffer^AVP^ cells were transiently transfected to express the red fluorescent genetically encoded calcium indicator R-GECO (GenScript, Piscataway, NJ, USA) with Fugene HD reagent (Promega, Madison, WI, USA). Sniffer^AVP^ cells were resuspended in standard aCSF (in mM): 119NaCl, 2.5 KCl, 1 MgSO 4, 26 NaHCO3, 1.25 NaH2 PO4, 20 D-glucose, 0.4 ascorbic acid, 2 CaCl2 and 2 pyruvic acid; pH 7.3; 300−305 mOsm) with trypsin (0.05%). Sniffer^AVP^ biosensors cells were transferred via pipette directly onto the SON of coronal brain slices submerged in aCSF in a perfusion-capable chamber of an upright microscope. For anatomical precision, the optic tract was used as a landmark. Before all recordings, eGFP-AVP neurons within the SON were observed in the field of view. An example of a SON slice containing both eGFP-AVP neurons and the Sniffer^AVP^ biosensors is shown in ***Fig.5***. After allowing the Sniffer^AVP^ biosensors to adhere to the slice for 5 minutes, aCSF superfusion was resumed to wash off any unattached Sniffer^AVP^ from the slice before proceeding with imaging. Experiments were restricted to preparations that had at least five fluorescently visible sniffer cells in the field (∼10 on average). To record the calcium-induced fluorescence changes of the Sniffer^AVP^ biosensors, images were captured using a Dragonfly 200 confocal scanning system and an iXON EMCCD camera (Andor Systems, Belfast, UK), at a rate of 1 Hz, with piezo-driven z-series to maximize biosensor counts per trial. Slices were constantly superfused with aCSF at 32°C at ∼2 ml/min. Imaging data were analyzed using ImageJ software (NIH). All data were background-subtracted. For quantitative measurements, fractional fluorescence (F/F0) was determined by dividing the fluorescence intensity (F) within a region of interest by the baseline fluorescence value (F0), which was determined from 30 frames preceding stimulation. Peak sniffer Ca^2+^ amplitude was the maximum F/F0 achieved following UV laser uncaging of MNI-caged-L-Glutamate (25 µM) in the presence of DNQX (10 µM) to restrict activation of AMPA receptors. As shown in ***Fig.5***, glutamate uncaging resulted in a robust burst firing in AVP neurons, which we previously shown to efficiently evoke local release of AVP within this SON ^34,35^. The area under the curve (F/F0∗s) was calculated by integrating the response over its duration. Response rates represent the number of biosensors that responded to the stimulation, divided by the total number of biosensors present in the field of view. Sniffer^AVP^ biosensors that showed intrinsic oscillatory calcium activity were excluded from the analysis. To better display changes in fluorescence levels, images were pseudo-colored using ImageJ.

### Statistical analysis

All statistical analyses were performed using GraphPad Prism 10 (GraphPad Software, California, USA). The significance of differences was determined using paired or unpaired t-tests (two-sided in all cases), one-way or two-way repeated-measures (RM) or normal analysis of variance (ANOVA) with the post hoc Bonferroni’s multiple comparisons test. Results are expressed as means ± standard deviation (SD). Results were considered to be statistically significant if p < 0.05 and are presented as ^∗^ for p < 0.05, ^∗∗^ for p < 0.01, and ^∗∗∗^ for p < 0.0001 in the respective figures.

## RESULTS

### Sustained hypoxic milieu in the SON of HF rats

A schematic of the 2P *in vivo* hypothalamic preparation is shown in ***Fig.1A***. Similar to our previous angioarchitecture characterization^6^, the SON in HF rats was readily identified by the presence of eGFP-AVP-expressing neurons which reside within the dense capillary network (labelled with i.v. injection of rhodamine 70kDa dextran dye, ***Fig.1B***). This dense capillary network is located medially to the bifurcation of the internal carotid artery (ICA), one of the main pial vessels supplying the SON^36^ and the branching of the middle cerebral artery (MCA) from the ICA (***Fig.1B***). As previously described, parenchymal arterioles (PA) originated near the most caudal aspect of the SON, which subsequently branched into smaller arterioles, forming this dense capillary network. We first determined whether, in sham vs HF, baseline vessel diameter differed across main SON arterioles (i.e., PA, ICA, MCA). *In vivo* 2-photon imaging of the SON vasculature revealed no difference in the different vessel types (i.e., PA, ICA and MCA) between sham and HF rats (F= 3.2, P= 0.08, two-way ANOVA, n=6 of each vessels/6 rats in each group) (***Fig.1C***). However, a significant decrease in basal blood flow was observed in PA of HF compared to sham rats (Sham: 30.01 ± 1.78 µm/ms; HF: 20.27 ± 3.18 µm/ms; n= 3 in sham and 4 in HF; p<0.01; unpaired t-test) **(*Fig. 1D*)**. We next assessed whether SON parenchymal pO2 saturation level was altered during HF. To this end, we placed an optical fluorescence probe (250 mm tip diameter; OxyLite system; Oxford Optronix) in the SON parenchyma, which allows real-time recording of parenchymal pO2 level ^6^. We found that SON pO2 saturation was lowered in HF compared to sham rats (sham: 22.87 ± 4.16 mmHg; HF: 16.66 ± 2.73 mmHg; n=6; p<0.05; unpaired t-test) **(*Fig. 1E*)**. Taken together, these results support sustained tissue hypoxia within the SON of HF rats at baseline conditions.

### Reversed polarity of the salt-evoked NVC in HF rats

Our previous work demonstrated that a systemic salt challenge, a potent stimulation of SON AVP neuronal activity, evoked a vasoconstriction of SON parenchymal arterioles (i.e., inversed NVC response, iNVC) that was mediated by somatodendritic release of AVP ^6^. Given that AVP expression and release are both enhanced in HF rats ^17,37,38^, we hypothesized a amplified iNVC response in this experimental group. To test this hypothesis, a group of sham and HF rats were subjected to the same salt challenge (2M NaCl, 2ml/hour, i.v. infusion for 30 min), while continuous PA diameter measurements were obtained under 2P imaging. As previously reported, we found that this salt challenge evoked a change in plasma osmolality of ∼40 mOsm/L, with no differences observed between sham and HF (sham: 361 ± 0.81 mOsm/L; HF: 359 ± 1.41 mOsm/L, n=4 in sham and n=5 in HF: p>0.05; unpaired t-test) (see ***supplementary Table 1***). Representative 2P images of SON PAs before and during the salt challenge in sham and HF rats are shown in ***Fig.2A1-A5*** and ***Fig.2B1-B5***, respectively. Consistent with our previous findings obtained in wild-type rats, sham rats in this study showed vasoconstriction of SON PA in response to acute salt loading (one-way RM ANOVA; F= 37.5, p< 0.001; n=6 vessels/6 rats) (***Fig.2C***). In contrast, the same procedure in HF rat evoked a pronounced vasodilation of SON PAs (one-way RM ANOVA; F= 102.8 p< 0.0001; n=6 vessels/6 rats) (***Fig.2C***). No noticeable differences in the onset of the vasoconstriction or vasodilation were observed, in both cases starting at ∼ 10 mins after the salt infusion was initiated. Furthermore, vascular responses persisted in both groups throughout the salt challenge.

**Figure 2.**
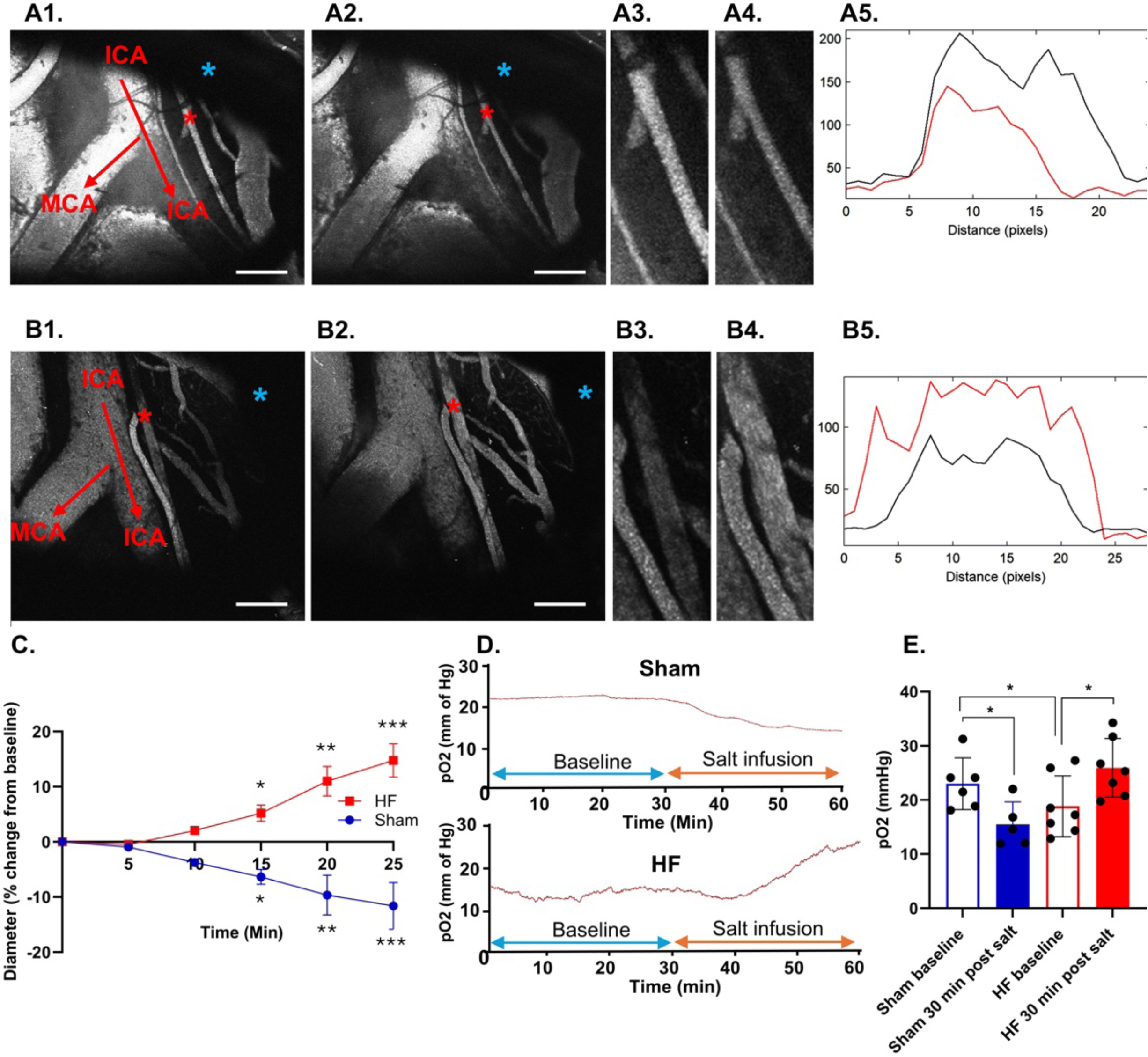
The polarity of the salt-evoked NVC response is reversed in HF rats. (**A**) Low magnification *in vivo* two-photon images of the SON following rhodamine 70 kDa administration (IV) before (A1) and during (A2) the salt challenge (2M NaCl, 1ml, 2ml/hour) in a representative sham rat. The blue asterisk indicates the shadow cast by the microdialysis probe. A3 and A4 represent the magnified image of the PA (demarcated as a red asterisk) in A1 and A2, respectively. In A5, the profiles of image brightness along the cross-section line of the PA vessels shown in A3 (before, black) and during (A4, red) the salt challenge are shown. Note the reduced PA diameter (vasoconstriction) during the salt challenge. (**B**) Low magnification *in vivo* two-photon images of the SON following rhodamine 70 kDa administration (IV) before (B1) and during (B2) the salt challenge (2M NaCl, 1ml, 2ml/hour) in a representative HF rat. B3 and B4 represent the magnified image of the PA (demarcated as a red asterisk) in B1 and B2, respectively. In B5, the profiles of image brightness along the cross-section line of the PA vessels, as shown in B3 (before, black) and during (B4, red) the salt challenge, are displayed. Note the increased PA diameter (vasodilation) during the salt challenge. (**C**) Summary plot of the mean percent change in PA diameter as a function of time during the salt challenge in sham (blue) and HF (red) rats. Note the significant vasoconstriction evoked in sham rats (blue, F= 37.5, p< 0.001, one-way RM ANOVA) and the vasodilation evoked in HF rats (red, F= 102.8, p < 0.0001, one-way RM ANOVA, n=6 vessels/6 rats in each experimental group). (**D**) Representative raw traces showing changes in SON tissue partial pressure of oxygen (pO2) before and during the salt infusion in a representative sham (upper) and HF (lower) rat. (**E**) Bar graph summarizing the mean SON pO2 before and 30 min after the salt challenge in sham (blue) and HF (red) rats. n=6 rats in each group. Scale bar=200mm. Error bars represent SEM. *p< 0.05, one-way ANOVA, Bonferroni’s post hoc test.

To determine if the salt-evoked NVC responses impacted the pO_2_ within the SON, we measured pO_2_ saturation levels during the salt challenge in both experimental groups. Representative traces showing the change in pO_2_ in the SON of a sham and HF rats are shown in ***Fig.2D*.** Consistent with the vasoconstriction response evoked in sham rats, the acute salt challenge significantly reduced SON pO_2_ saturation SON in sham rats (baseline: 22.99 ± 4.35 mmHg; after salt loading: 15.47± 3.75 mmHg; n=6; p<0.05; unpaired t-test) (***Fig.2E***). Conversely, but consistent with the vasodilatory response in HF rats, the acute salt challenge significantly increased the pO_2_ saturation in the SON of HF rats (baseline: 18.84 ± 5.19 mmHg; after salt loading: 25.93 ± 5.01 mmHg; n=7; p<0.05; Bonferroni’s *post-hoc* test), normalizing the baseline hypoxic condition, to similar baseline levels observed in sham rats (***Fig.2E***). Taken together, these results support a change in the polarity of the vascular response (from vasoconstriction to vasodilation) in the salt-evoked NVC response in HF rats, which effectively resulted in an increase and normalization of the pO_2_ saturation level within the SON.

### The salt-evoked vasodilation in HF rats is mediated by adenosine acting on A2AR receptors

Previously, we demonstrated that the salt-evoked vasoconstriction in the SON was mediated via local dendritic release of AVP ^6^. In HF rats, microdialysis of AVP directly onto the SON (1 mM, 2.2ml/hour for 5 min) also evoked a progressive vasoconstriction of SON PAs (one-way RM ANOVA, F= 20.8, p<0.0001, n=5 vessels/5 rats) (**Fig.3A-C**). These results suggest that, in HF rats, AVP is not the main signal mediating the salt-evoked vasodilatory response. One possibility is that the high salt stimulus in HF leads to the release of a vasodilatory signal that overrides the AVP-induced vasoconstriction observed in healthy rats. A potential candidate is adenosine. Adenosine, acting on adenosine A_2A_ receptors (A_2A_R), causes potent vasodilation ^39–42^ and can contribute to NVC response in the somatosensory cortex ^43,44^ and retina ^45^. To investigate whether a similar signaling cascade could contribute to the salt-evoked vasodilation in HF rats, we first tested, as a proof of concept, whether microdialysis of CGS 21680, a selective A_2A_R agonist, directly onto the SON of HF rats (1 mM, 2.2ml/hour for 5 min) induced a vasodilatory response. As shown in **Fig.3B**, we found that CGS 21680 caused a progressive vasodilation of SON PAs in HF rats (one-way RM ANOVA, F= 21.0, n=4 vessels/4 rats; p<0.01). These results support a potential contribution of endogenously produced adenosine to the salt-evoked vasodilatory response in HF rats. To test this, we performed a separate set of experiments in which HF rats were subjected to a systemic salt challenge (2M NaCl, IV, 1ml/hour) while ZM 241385, a selective A_2A_R antagonist, was microdialyzed directly into the SON (1mM, 0.6ml/hour). As shown in ***Fig.4* *A1-A5***, we found that local delivery of the A_2A_R antagonist in the SON not only prevented the salt-induced vasodilation, but in fact, it unmasked an underlying vasoconstriction (one-way RM ANOVA; F= 35.9, p < 0.001; n=6 vessels/6 rats) (***Fig.4D***). Importantly, this unmasked vasoconstriction was abolished when the A_2A_R antagonist was co-infused into the SON with the AVP V1aR antagonist (β-mercapto-β,-cyclopentamethylenepropionyl^1^, [O-me-Tyr^2^, Arg^8^]-AVP, 200 μM) (***Fig.4B1-B5 and 4D***) (one-way RM ANOVA; F= 1.4 p=0.6; n=4 vessels/4 rats). Together, these results support the notion that in response to a salt challenge, in HF rats, adenosine via A_2A_Rs, overrides the AVP-induced vasoconstriction mediated via V1aR activation.

**Figure 3.**
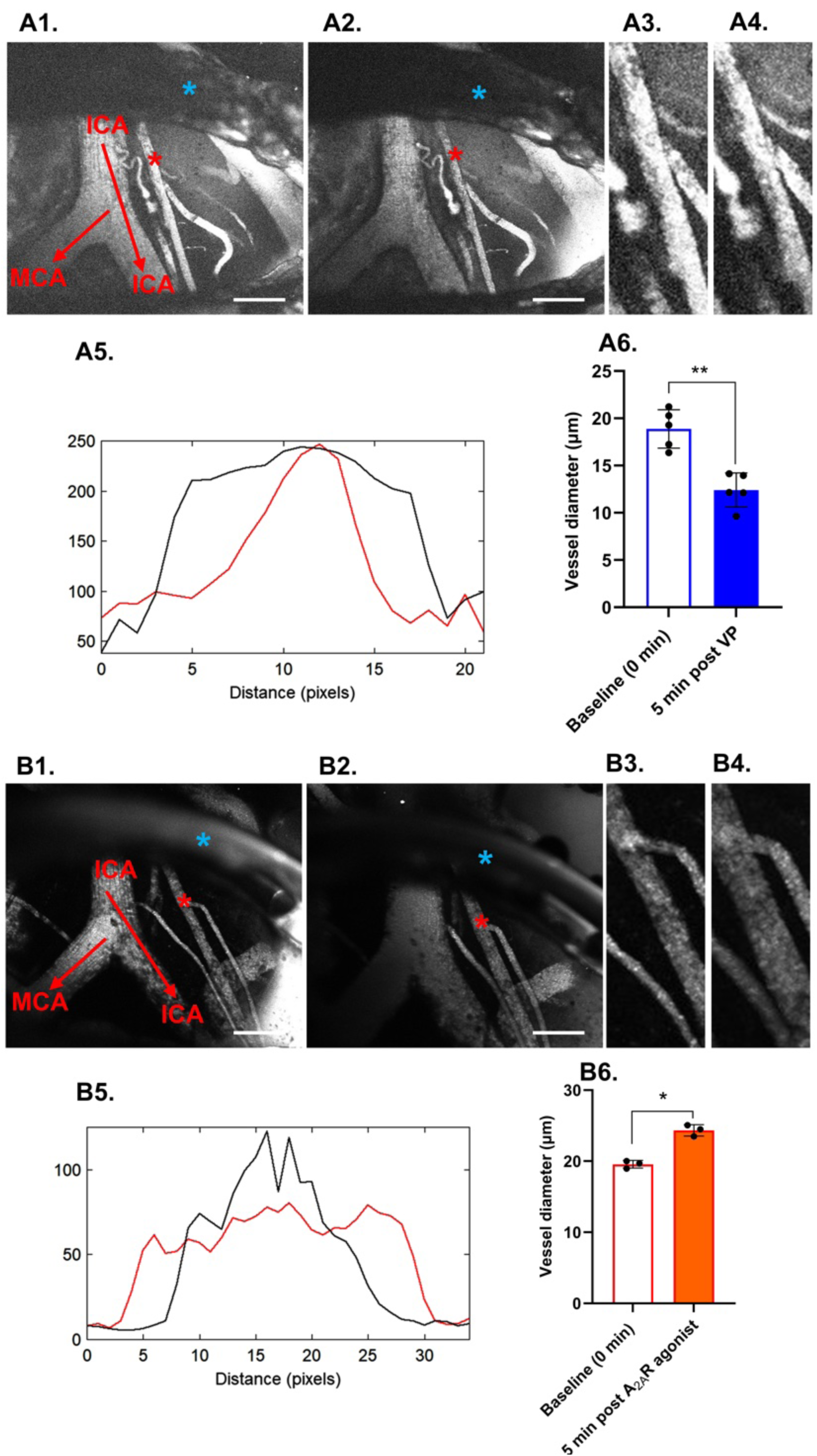
Vasopressin (AVP) and adenosine evoke vasoconstriction and vasodilation, respectively, in SON parenchymal arterioles (PA) in HF rats. **(A)** Low magnification *in vivo* two-photon images of the SON following rhodamine 70 kDa administration (IV) before (A1) and during (A2) topical infusion of AVP (1 mM, 2.2ml/hour) through a microdialysis probe positioned on top of the SON in a representative HF rat (the blue asterisk indicates the shadow cast by the microdialysis probe). A3 and A4 represent the magnified image of the PA (demarcated as a red asterisk) in A1 and A2, respectively. In A5, the profiles of image brightness along the cross-section line of the PA vessels shown in A3 (before, black) and in A4 (during, red), the AVP infusion, are displayed. Note the reduced PA diameter (vasoconstriction) evoked by AVP. A6, Bar graph showing the mean changes in PA diameter after the local infusion of AVP. n=5 vessels/5 rats; **(B)** Low magnification *in vivo* two-photon images of the SON following rhodamine 70 kDa administration (IV) before (B1) and during (B2) topical infusion of the adenosine A_2A_ receptor agonist CGS 21680 (1 mM, 2.2ml/hour) through a microdialysis probe positioned on top of the SON in a representative HF rat (the blue asterisk indicates the shadow cast by the microdialysis probe). B3 and B4 represent the magnified image of the PA (demarcated as a red asterisk) in B1 and B2, respectively. In B5, the profiles of image brightness along the cross-section line of the PA vessels shown in B3 (before, black) and in B4 (during, red), the CGS 21680 infusion, are displayed. Note the increased PA diameter (vasodilation) evoked by CGS 21680. B6, Bar graph showing the mean changes in PA diameter after the local infusion of CGS 21680. n=3 vessels/3 rats; Scale bar=200mm. Error bars represent SEM. * p<0.05 and **p< 0.01 paired t-test.

**Figure 4:**
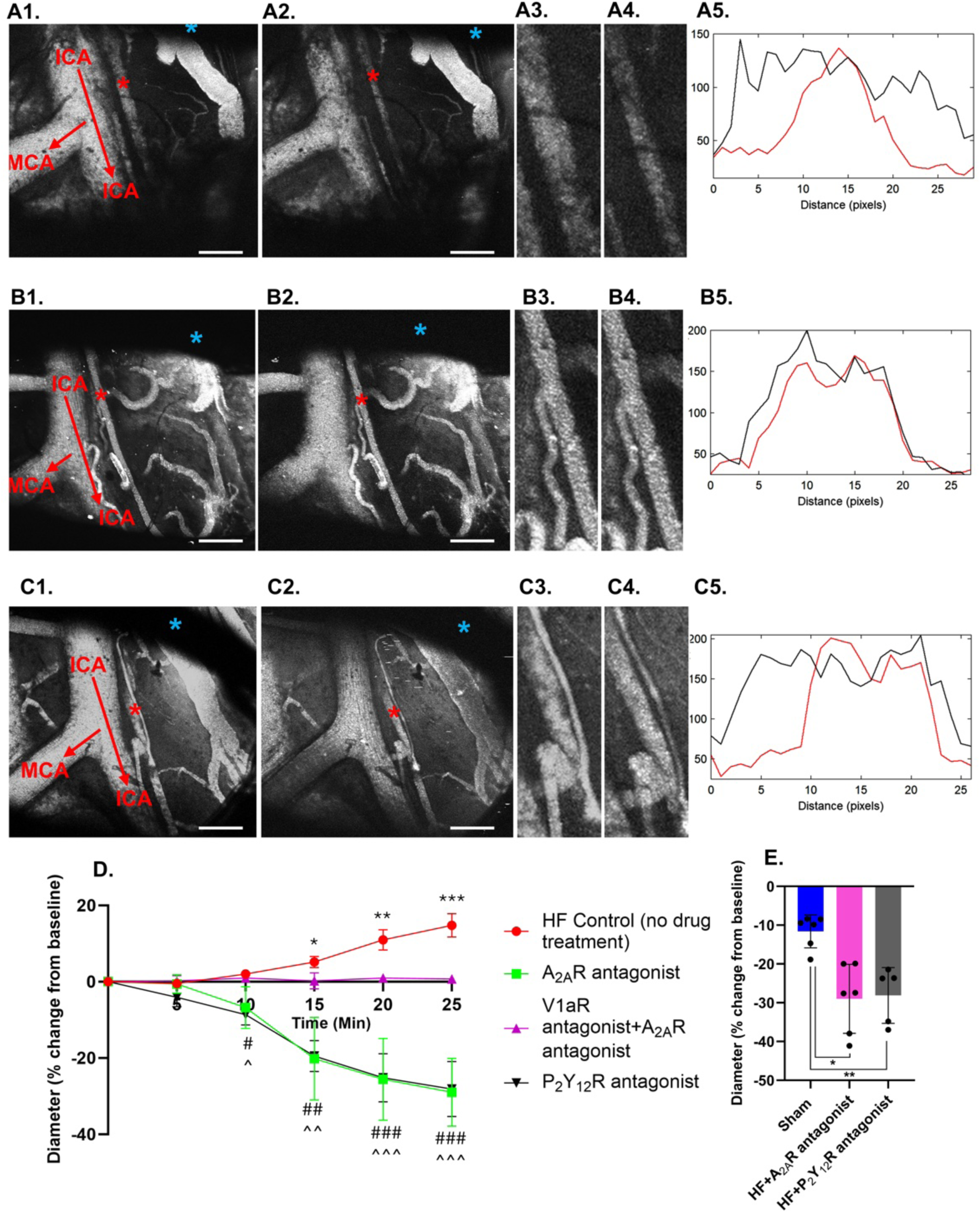
The salt-evoked vasodilatory response in HF rats is mediated by endogenously released adenosine acting on A_2A_ receptors. (**A**) Low magnification *in vivo* two-photon images of the SON following rhodamine 70 kDa administration (IV) before (A1) and during (A2) the salt challenge (2M NaCl, 1ml, 2ml/hour) in the presence of microdialysed ZM 241385 (a selective A_2A_R antagonist,1mM, 0.6ml/hour) in a representative HF rat. The blue asterisk indicates the shadow cast by the microdialysis probe. A3 and A4 represent the magnified image of the PA (demarcated as a red asterisk) in A1 and A2, respectively. In A5, the profiles of image brightness along the cross-section line of the PA vessels shown in A3 (before, black) and during (A4, red) the salt challenge, are displayed. Note the reduced PA diameter (vasoconstriction) during the salt challenge in the presence of the A_2A_R blocker. (**B**) Low magnification *in vivo* two-photon images of the SON following rhodamine 70 kDa administration (IV) before (B1) and during (B2) the salt challenge in the presence of co-microdialysed (0.6ml/hour) ZM 241385 (1 µM) plus β-mercapto-β,-cyclopentamethylenepropionyl (a V1a receptor antagonist, 200 mM) in another representative HF rat. B3 and B4 represent the magnified image of the PA (demarcated as a red asterisk) in B1 and B2, respectively. In B5, the profiles of image brightness along the cross-section line of the PA vessels shown in B3 (before, black) and during (B4, red) the salt challenge, are displayed. Note lack of changes PA diameter during the salt challenge in the presence of both A_2A_R and V1a receptor blockers. (**C**) Low magnification *in vivo* two-photon images of the SON following rhodamine 70 kDa administration (IV) before (C1) and during (C2) the salt challenge in the presence of microdialysed PSB 0739 (a selective purinergic receptor P_2_Y_12_ antagonist, 1 µM 0.6ml/hour) in another representative HF rat. C3 and C4 represent the magnified image of the PA (demarcated as a red asterisk) in C1 and C2, respectively. In C5, the profiles of image brightness along the cross-section line of the PA vessels shown in C3 (before, black) and during (C4, red) the salt challenge, are displayed. Note the reduced PA diameter (vasoconstriction) during the salt challenge in the presence of the P_2_Y_12_ blocker. **D**, Summary plot of the mean percent change in PA diameter as a function of time during the salt challenge in HF rats in the control conditions (red), or in the presence of microdialysed A_2A_R antagonist (green), P_2_Y_12_R antagonist (black), or a combination of a V1aR antagonist with the A_2A_R antagonist (purple). n=5 vessels/5 rats in each group. *p<0.05, **p<0.01, ***p<0.001, control group. #p<0.05, ##p<0.01, ###p<0.001, for A_2A_R antagonist group, #p<0.05, ##p<0.01, ###p<0.001, P_2_Y_12_R antagonist group, one-way ANOVA followed by Bonferroni’s post-hoc test, vs baseline. **E,** Bar graph showing the mean peak magnitude of the salt-evoked vasoconstriction in sham rats (blue, n=6 vessels/6 rats), and HF rats in the presence of the A_2A_R antagonist (pink, n=6 vessels/6 rats) and the P_2_Y_12_R antagonist (black, n=5 vessels/5 rats). Scale bar=200mm. Error bars represent SEM. **p<0.01, one-way ANOVA, Bonferroni’s post-hoc test.

**Figure 5:**
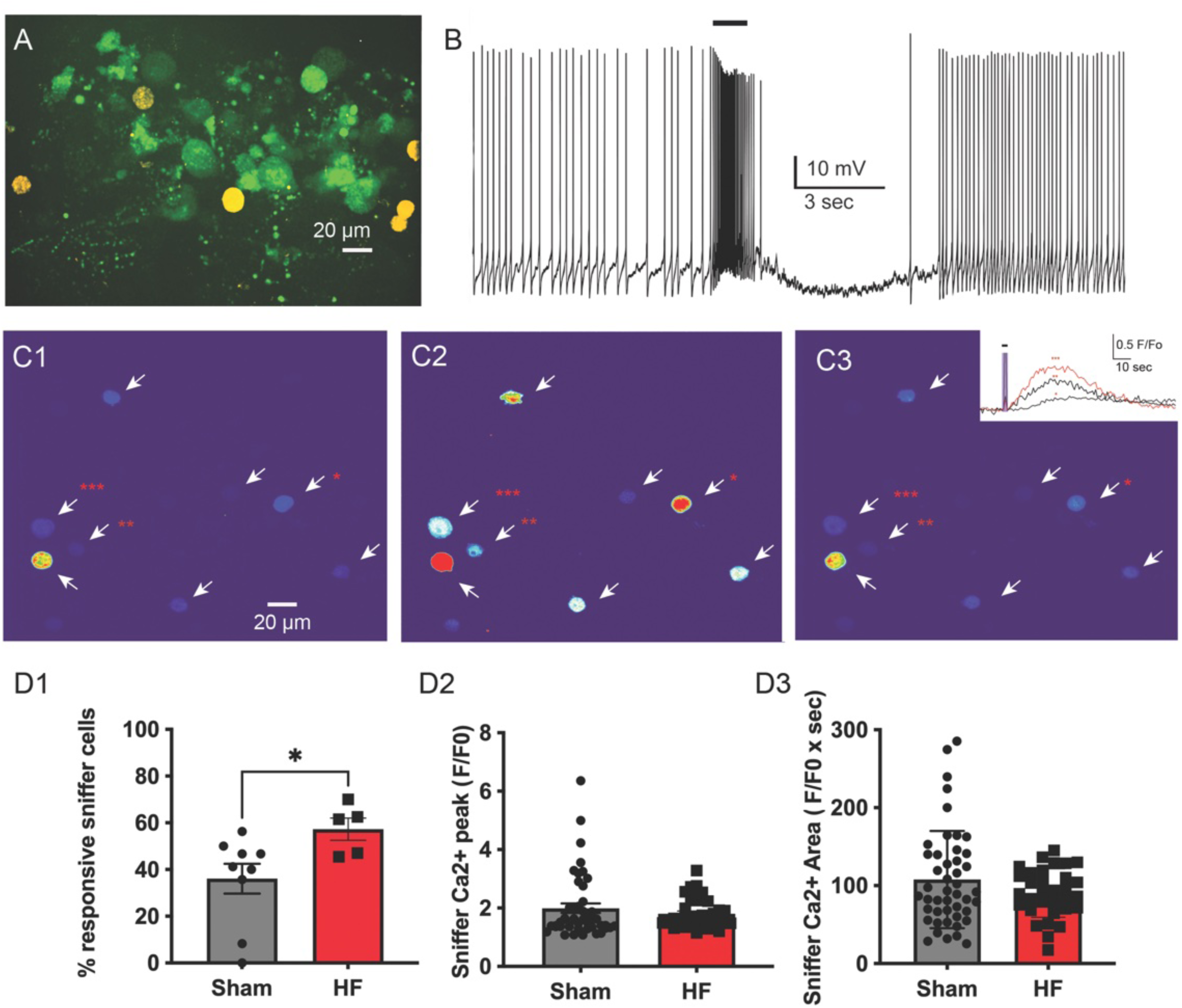
Enhanced activity-dependent local release of AAVP within the SON of HF rats. (**A**) Representative image of the SON in a hypothalamic slice containing eGFP-AAVP neurons (green) and the sniffer-AAVP biosensor cells (yellow) plated on the SON. (**B**) Representative example of a patched AAVP-GFP neuron showing a robust burst of action potentials evoked by focal glutamate uncaging (UV light, 2 seconds, black bar). (**C**) Pseudocolour Ca^2+^ images showing responsive Sniffer-AAVP cells (arrows) at baseline (C1), immediately after stimulation (C2), and the recovery after stimulation(C3) following glutamate uncaging in a representative slice obtained from an HF rat. The inset in C2 shows corresponding Ca^2+^ traces of the sniffer-AAVP cells indicated with the corresponding asterisks. (**D**) Bar graphs showing mean data of the percentage of responsive sniffer-AAVP cells (D1), Ca^2+^ peak (D2) and Ca^2+^ area (D3) in sham and HF rats. *p< 0.05 (unpaired t-test).

### Microglia contribute to adenosine-mediated, salt-evoked vasodilatory responses in HF rats

In the central nervous system (CNS), microglia constitute an important source of adenosine ^46,47^, and recent studies indicate that microglia actively regulate CBF via an adenosine signaling mechanism ^48,49^. Moreover, the purinergic receptor P_2_Y_12_ is exclusively expressed on microglia^50,51^, activation of which plays a pivotal role in microglial activation and the release of signals in response to various stimuli ^52–54^. To investigate whether microglia contribute to the salt-evoked vasodilation of SON PA in HF rats, we repeated a systemic salt challenge in a separate set of experiments while PSB0739, a selective P2Y12 purinergic receptor antagonist, was microdialyzed directly into the SON (1 µM, 0.6ml/hour). Similarly to what we observed with the A_2A_R antagonist, PSB0739 not only prevented the salt-evoked vasodilation but also unmasked an underlying vasoconstriction in HF rats (one-way RM ANOVA; F= 77.8, p < 0.001; n=5 vessels/5 rats) (***Fig.4C1-C5 and 4D***). Taken together, these results suggest that microglia constitute a potential source of adenosine, contributing to the salt-evoked vasodilation of SON PA in HF rats.

Together, the above results indicate that in HF rats, salt stimulates the release of both AVP and adenosine within the SON, having opposing effects on the local microvasculature. To gain insights into whether salt activates these two signaling mechanisms in parallel, or whether they act in series (e.g., salt evokes the release of AVP which then induces the release of adenosine), we repeated the systemic salt challenge in a separate set of experiments in HF rats in the continuous presence of the V1aR antagonist (microdialysis of 200 μM, 0.6ml/hour). In this condition, we found no changes in vascular diameter (neither vasoconstriction nor vasodilation) in response to the salt challenge (One-way RM ANOVA; F= 1.68, p= 0.4; n = 5 vessels/5 rats ***supplementary Fig: 1***). Taken together, these results suggest that salt-induced vasodilation of SON PA in HF rats, while mediated by adenosine acting on A_2A_Rs, is dependent on an endogenous AVP-mediated signaling cascade.

### An increased activity-dependent somatodendritic release of AVP likely contributes to an enhanced AVP-mediated vasoconstriction (iNVC) in HF rats

Notably, we found that the magnitude of the “masked” salt-evoked, AVP-mediated vasoconstriction in HF rats (i.e., in the presence of the A_2A_R or the P_2_Y_12_R antagonists) was significantly larger than that evoked in sham rats (HF-A_2A_R antagonist: -29.0 ± 5.37%, HF-P2Y12R antagonist:-28.12 ± 6.44%; sham: -11.62 ± 3.67%; n=6 vessels/6 rats in each group; One way ANOVA, F= 11.3, p<0.001; unpaired t-test) **(*Fig. 4E*)**. These results indicate that even though the salt-evoked adenosine-mediated vasodilation overrides the AVP-mediated vasoconstriction in HF rats, somatodendritic release of AVP may still be enhanced in this condition. To test whether this was the case, we used Sniffer^AVP^ biosensor cells (Methods), an approach we and others recently demonstrated to detect endogenous release of neuropeptides with extremely high sensitivity and spatiotemporal resolution ^55–57^. Sniffer^AVP^ biosensors were plated into *ex vivo* slices containing the SON, obtained from either sham or HF rats (n=4 and 5 sham and HF rats, respectively). Photolysis of caged-glutamate was used to stimulate NMDA receptor-evoked firing and somatodendritic release of AVP within the SON (see Methods), an approach we previously demonstrated to be highly efficient in evoking this local release modality ^34^. To validate the photolysis approach, we first obtained patch clamp recordings from identified AVP neurons under control conditions, demonstrating that our laser “uncaging” protocol worked as intended. As shown in ***Fig.5***, laser photolysis of caged glutamate (UV flash, 2 seconds) induced a robust burst of action potential in patched AVP neurons (mean number of action potentials, 38.3 ± 6.4, mean interspike frequency: 22.8 ± 3.1 Hz (significantly higher than baseline frequency of 3.4 ± 0.9 Hz, p< 0.0001, paired t test, n=8). As previously reported, we found that NMDA-evoked firing in AVP neurons resulted in local AVP release, as shown by the responsiveness of Sniffer^AVP^ biosensors (i.e., increased in Ca^2+^ fluorescence). Importantly, we found that the percentage of responsive Sniffer^AVP^ biosensors was significantly higher in slices obtained from HF (∼57%, 38/67) compared to sham rats (∼37%, 46/124, p< 0.05, unpaired t-test, ***Fig.5***). Conversely, the peak and area of the Ca^2+^ response in the responsive Sniffer^AVP^ biosensors was similar between the two experimental groups (p= 0.7 and 0.1, for peak and area, respectively, unpaired t-test, n= 46 and 38 Sniffer^AVP^ biosensors from sham and HF rats, respectively, ***Fig.5***). Together, these results support an increased activity-dependent somatodendritic release of AVP in HF rats.

## DISCUSSION

Neurovascular coupling (NVC) is a critical brain process that ensures increased cerebral blood flow (CBF) in response to neuronal activity ^1,2^. Recently, an impaired NVC response in the cortex has been reported both in animal models ^13^ and human patients ^8^ with heart failure (HF), a pathophysiological mechanism proposed to contribute to cognitive impairment and dementia in this prevalent cardiovascular disease. Most existing studies on NVC, however, are limited mainly to superficial dorsal brain areas, particularly the cortex, and focus on responses to specialized sensory inputs. In these cases, neuronal-activity-evoked CBF changes occur rapidly (<2 s) and in a spatially restricted manner ^4^. Recently, we addressed this limitation by implementing a novel *in vivo*, real-time two-photon experimental approach to assess NVC for the first time in deep brain hypothalamic nuclei, including the SON ^6,58^. In this recent work, we demonstrated that as opposed to the conventional functional hyperemic response previously described, increased activity of SON AVP neurons via a systemic salt challenge evoked parenchymal arteriole vasoconstriction (hence termed as inverse NVC (iNVC)) and a progressive decrease in SON pO_2_. Moreover, we demonstrated that iNVC was mediated by local somatodendritic release of AVP within the SON ^6^.

The hypothalamus is a critical brain region for the generation of homeostatic neurohumoral responses ^59^. Moreover, a large body of evidence supports a hypothalamic contribution to exacerbated neurohumoral responses in HF, resulting in higher morbidity and mortality in this disease ^60–62^. Given that excess dietary salt triggers neurohumoral responses involving sympathoexcitation and the release of AVP ^63,64^, and that it worsens fluid retention and congestion in HF ^22^, we considered it imperative to determine if salt-evoked NVC responses were altered in HF, in a way that could contribute to the pathophysiology of this disease.

Nitric oxide (NO) is a major systemic and central vasodilator ^65^, and a blunted hypothalamic NO expression and function has been reported in HF rats ^17,20,21^. Conversely, the expression and release of AVP, a potent vasoconstrictor, are elevated in HF rats ^16,19^. Based on the imbalance of these two major vasoactive modulators favoring vasoconstriction in HF, and the fact that we showed salt-evoked iNVC to be mediated by AVP ^6^, we hypothesized that exacerbated salt-evoked vasoconstrictions would occur in the SON of HF rats, leading to a more pronounced hypoxic milieu in this condition.

Surprisingly, we observed quite the opposite response. Our main results can be summarized as follows: 1) under baseline conditions, HF rats exhibit lower levels of pO_2_ in SON; 2) Despite this initial basal SON hypoxic condition, a salt challenge induced parenchymal arteriolar vasodilatation and a progressive increase in pO_2_, restoring it to comparable levels as those observed in Sham rats; 3) The salt-evoked vasodilation in HF was mediated by adenosine, acting on adenosine A_2A_ receptors, of a putative microglial source; 4) A masked AVP-mediated vasoconstriction was still observed, which was even of a larger magnitude to that observed in sham rats, and was likely due to an enhanced activity-dependent somatodendritic release of AVP, as shown by our biosensor sniffer cell studies.

### A Shift in the balance of vasoactive signals during HF impacts the polarity of the salt-evoked NVC

Using both *in vivo* and *ex vivo* approaches, we previously demonstrated that the neurohormone AVP regulates vascular tone and reactivity of parenchymal arterioles in the SON by inducing vasoconstriction through its action on V1a receptors on vascular smooth muscle cells ^66^. This included a vasoconstrictive NVC response to a systemic salt challenge ^6^. Notably, we also demonstrated that when AVP signaling was pharmacologically blocked, a previously masked vasodilatory response was revealed ^6^, which was likely mediated by nitric oxide (NO) ^66^. This suggests that a salt challenge activates a mix of vasodilatory and vasoconstrictor signals, and that under normal conditions, the AVP-mediated signal predominates, resulting in an overall vasoconstrictive response. Nonetheless, the masked vasodilation may function to limit the duration and magnitude of this vasoconstrictive response, preventing excessive vasoconstriction during the sustained salt challenge, and ensuring dynamic vascular homeostasis.

A significant finding of the present study is a drastic change in the polarity of the salt-evoked NVC response in rats with HF. Rather than vasoconstriction being the dominant response, we found that the same homeostatic challenge (i.e., salt) resulted in a vasodilatory response of parenchymal arterioles in the SON. This vasodilation was primarily mediated by adenosine acting on A_2A_ receptors, which are known to be preferentially expressed on the vasculature ^67^ and to promote vascular smooth muscle relaxation ^68^. Interestingly, the adenosine-induced vasodilation in HF rats masks an underlying AVP-mediated vasoconstriction, which became evident only when A_2A_ receptor signaling was topically blocked within the SON. Thus, similar to control conditions, the salt challenge in HF rats resulted in the release of competing vasoactive signals. However, in this condition, vasodilation predominated over vasoconstriction.

Despite the predominant adenosine/A_2A_-mediated vasodilation, we found that the magnitude of the masked AVP-mediated vasoconstriction was larger in HF rats compared to sham rats, suggesting an enhanced activity-dependent somatodendritic release of AVP in the former. We and others have previously shown that the excitability of AVP neurons is enhanced in HF rat ^18,69,70^, which leads to increased circulating levels of the hormone ^19,71^. However, whether an augmented systemic release is accompanied by central somatodendritic release of AVP has not been tested before. Our studies using a biosensor sniffer cell support the notion that activity-dependent central release of AVP is also enhanced in HF, a mechanism likely contributing to the more pronounced but “masked” AVP-mediated vasoconstriction in this condition. Still, whether this is directly due to the increased excitability of AVP neurons or other factors influencing somatodendritic release (e.g., intracellular Ca^2+^ homeostasis, increased AVP dendritic cargo) remains to be determined.

### Microglia as a putative source of adenosine mediating salt-evoked vasodilation in HF rats

Accumulating evidence supports that adenosine and its receptors, particularly the A2 subtypes (A_2A_ and A_2B_), contribute to CBF regulation ^72–74^, including NVC responses ^48,75^. For example, adenosine release in response to hypoxia leads to cortical pial vessel dilation and increased cortical CBF, effects prevented by A_2A_ receptor blockade ^76^. Moreover, a reduced CBF was observed in A_2A_ receptor knock-out mice ^77^. Finally, adenosine receptors were also implicated in mediating NVC response in the barrel cortex in response to whisker stimulation ^78^. Together, these findings underscore the crucial role of adenosine, particularly through A_2A_ receptor signaling, in regulating CBF and NVC responses.

Adenosine is synthesized and released through multiple pathways, primarily from the breakdown of ATP ^79^. Microglia, the resident immune cells of the CNS, have recently emerged as a significant source of extracellular adenosine. Upon activation by various stimuli such as inflammation, hypoxia, or neuronal activity, microglia release ATP, which is rapidly converted to adenosine by the ectonucleotidases CD39 and CD73 expressed on microglial surface ^80,81^. Notably, recent studies support the role of microglia in mediating NVC vasodilatory responses ^48,82,83^. In the present study, we demonstrate that, in addition to A_2A_ receptor blockade, the salt-evoked vasodilation in HF rats was also prevented by selective blockade of P_2_Y_12_ receptors. These are G-protein-coupled receptors, which are predominantly expressed on the surface of microglia in the CNS ^84,85^. P_2_Y_12_ receptors are essential for microglial functioning and their response to a variety of signals, including those from neuronal origins ^85–89^. Moreover, a recent study has shown that microglia-mediated NVC responses in the cortex are dependent on the normal function of their P_2_Y_12_ receptors, ^82^. Based on this, we found it reasonable to speculate that microglia constitute an essential source of adenosine contributing to the salt-evoked vasodilatory responses in HF rats.

A contribution of microglia to the salt-evoked vasodilatory response is reasonable in the context of HF. In this sense, we and others demonstrated that neuroinflammation, including increased microglial activation, constitutes a major pathophysiological feature in the hypothalamus and other brain regions during HF ^90–94^. Moreover, we recently reported an increased proportion of vessel-associated microglia in several brain regions of HF rats ^95^. Finally, in conditions of hypoperfusion/hypoxia, vessel-associated microglia increased their process motility and contributed to adaptive responses to cortical CBF reduction, actions that were also shown to be dependent on P_2_Y_12_ receptors ^82^. This is mechanistically relevant given that in the present study, we report a condition of sustained, basal hypoperfusion/hypoxia in the SON of HF rats. Together, these results suggest that the structural remodeling in the microglia/microvascular space during HF, along with the functional priming induced by the hypoxic milieu, could act as factors that facilitate the ability of microglia in this condition to influence vascular dynamics and to dominate the polarity of the NVC response. Still, while our results strongly support microglia as a source of adenosine mediating the salt-induced vasodilation observed in HF, we can’t rule out other possible cellular sources, such as endothelial and/or neuronal ones.

### AVP is the likely signal stimulating microglia to release adenosine in HF

Our studies indicate that the acute salt challenge in HF rats leads to the release of both AVP and adenosine within the SON. However, whether these signaling mechanisms act in parallel or if one signal triggers the activation of the other one is not fully determined. One possibility is that, in addition to stimulating AVP neuronal activation and leading to AVP release, salt can independently act on microglia to release adenosine. Indeed, high salt has been shown to result in microglia activation and a neuroinflammatory response, affecting in turn various conditions, including multiple sclerosis symptoms ^96^, ischemia ^97^, and stress-coping responses ^98^. Alternatively, it is now well established that microglia express receptors for multiple neurotransmitters, including neuropeptides ^99^, and can thus sense and respond to local neuronal activity. For example, several studies have shown that oxytocin acting on microglia exerts an anti-inflammatory effect ^100–103^. More importantly, it was also shown that AVP can evoke Ca^2+^ responses in isolated microglial cells, an effect that was significantly increased after microglia were induced into a pro-inflammatory state ^104^. These results suggest that salt-evoked AVP release within the SON could be the signal triggering microglia to release adenosine. This is in fact supported by our results showing that blocking AVP V1a receptors within the SON in HF rats before the salt challenge prevented all vascular responses (i.e., the adenosine-mediated vasodilation and the “masked” AVP-mediated vasoconstrictions). Thus, while these results suggest that AVP is upstream of the adenosine signal, whether this involves AVP acting on V1a receptors expressed in microglia remains to be determined.

In summary, our results demonstrate that in HF, a condition as we show here, is characterized by a basal hypoperfusion/hypoxic state of the hypothalamic SON, the salt-evoked NVC response switches polarity, resulting in a predominant vasodilatory response, which tends to reverse the hypoperfusion state of the SON. These effects are mediated, at least in part, by adenosine of a microglial source acting on A_2a_ receptors. In agreement with recent studies in other brain regions ^49^ our results further support the notion that microglia can dynamically regulate CBF and NVC responses, particularly in conditions of brain hypoperfusion. Given that compromised CBF is a condition commonly observed in other cardiovascular diseases, including stroke, hypertension and vascular dementia, among others ^2,105,106^, our results could have a broader impact on cardiovascular pathophysiology.

## ACKNOWLEDGEMENTS

Grant support was provided by NIH R01HL162575 to JES and JAF; AHA 916907 to RKR. We also acknowledge support provided by the Center for Neuroinflammation and Cardiometabolic Diseases at GSU.

---

**Supplementary Figure 1: The Salt-evoked, adenosine-mediated vasodilation in HF rats is dependent on an intrinsic AVP signaling mechanism. A** Low magnification *in vivo* two-photon images of the SON following rhodamine 70 kDa administration (IV) before (A1) and during (A2) the salt challenge (2M NaCl, 1ml, 2ml/hour) in the presence of microdialysed V1a receptor blocker β-mercapto-β,-cyclopentamethylenepropionyl (200μM, 0.6ml/hour) in a representative HF rat. The blue asterisk indicates the shadow cast by the microdialysis probe. A3 and A4 represent the magnified image of the PA (demarcated as a red asterisk) in A1 and A2, respectively. **B**, The profiles of image brightness along the cross-section line of the PA vessels shown in A3 (before, black) and during (A4, red) the salt challenge are shown. Note the lack of changes PA diameter during the salt challenge in the presence of the V1aR blocker. **C**, Summary plot of the percent change in PA diameter as a function of time during the salt challenge in HF rats in the presence of microdialysed V1aR antagonist (green). n=5 vessels/5 rats in each condition; F= 0.76, p= 0.69, one way ANOVA. Scale bar=200μm. Error bars represent SEM.

**Supplemental Table 1.**
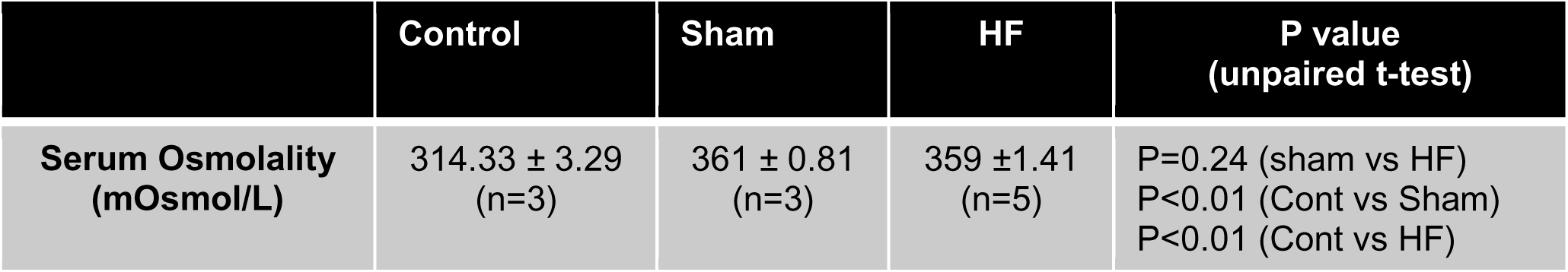
Summary of serum osmolality in control (no salt loading), sham and heart failure (HF) rats after salt loading.

|  | Control | Sham | HF | P value<br>(unpaired t-test) |
| --- | --- | --- | --- | --- |
| **Serum Osmolality**<br>**(mOsmol/L)** | 314.33 ± 3.29<br>(n=3) | 361 ± 0.81<br>(n=3) | 359 ± 1.41<br>(n=5) | P=0.24 (sham vs HF)<br>P<0.01 (Cont vs Sham)<br>P<0.01 (Cont vs HF) |

## References

1 Urban, A. et al. Understanding the neurovascular unit at multiple scales: advantages and limitations of multi-photon and functional ultrasound imaging. Advanced drug delivery reviews 119, 73–100 (2017).

2 Iadecola, C. The neurovascular unit coming of age: a journey through neurovascular coupling in health and disease. Neuron 96, 17–42 (2017).

3 Li, N. et al. High spatiotemporal resolution imaging of the neurovascular response to electrical stimulation of rat peripheral trigeminal nerve as revealed by in vivo temporal laser speckle contrast. J. Neurosci. Methods 176, 230–236 (2009).

4 Silva, A. C., Lee, S.-P., Iadecola, C. & Kim, S.-G. Early temporal characteristics of cerebral blood flow and deoxyhemoglobin changes during somatosensory stimulation. Journal of Cerebral Blood Flow & Metabolism 20, 201–206 (2000).

5 Meyer-Baese, L., Jaeger, D. & Keilholz, S. Neurovascular coupling: a review of spontaneous neocortical dynamics linking neuronal activity to hemodynamics and what we have learned from the rodent brain. J. Neurophysiol. 133, 644–660 (2025).

6 Roy, R. K. et al. Inverse neurovascular coupling contributes to positive feedback excitation of vasopressin neurons during a systemic homeostatic challenge. Cell reports 37 (2021).

7 Wareham, L. K. et al. Solving neurodegeneration: common mechanisms and strategies for new treatments. Mol. Neurodegener. 17, 23 (2022).

8 Aires, A. et al. Neurovascular coupling impairment in heart failure with reduction ejection fraction. Brain Sciences 10, 714 (2020).

9 Houben, A. J., Beljaars, J. H., Hofstra, L., Kroon, A. A. & LEEUW, P. W. D. Microvascular abnormalities in chronic heart failure: a cross-sectional analysis. Microcirculation 10, 471–478 (2003).

10 Youwakim, J., Vallerand, D. & Girouard, H. Neurovascular coupling in hypertension is impaired by IL-17A through oxidative stress. Int. J. Mol. Sci. 24, 3959 (2023).

11 Lefferts, W. K., DeBlois, J. P., Barreira, T. V. & Heffernan, K. S. Neurovascular coupling during cognitive activity in adults with controlled hypertension. J. Appl. Physiol. 125, 1906–1916 (2018).

12 Lin, W., Hao, Q., Rosengarten, B., Leung, W. & Wong, K. Impaired neurovascular coupling in ischaemic stroke patients with large or small vessel disease. Eur. J. Neurol. 18, 731–736 (2011).

13 Hoyer-Kimura, C. et al. PNA5, a novel mas receptor agonist, improves neurovascular and blood-brain-barrier function in a mouse model of vascular cognitive impairment and dementia. Aging and disease 15, 1927 (2024).

14 Lambrichts, S. M. et al. Vascular contribution to cognitive impairment in heart failure with preserved ejection fraction: TRPV4 and KLF2 as key mediators of neurovascular dysfunction in the ZSF1 model. bioRxiv, 2025.2001. 2008.631937 (2025).

15 Cohn, J. N. et al. Plasma norepinephrine as a guide to prognosis in patients with chronic congestive heart failure. N. Engl. J. Med. 311, 819–823 (1984).

16 Patel, K. P., Zhang, K., Kenney, M. J., Weiss, M. & Mayhan, W. G. Neuronal expression of Fos protein in the hypothalamus of rats with heart failure. Brain research 865, 27–34 (2000).

17 Reis, W. L., Biancardi, V. C., Zhou, Y. & Stern, J. E. A functional coupling between carbon monoxide and nitric oxide contributes to increased vasopressin neuronal activity in heart failure rats. Endocrinology 157, 2052–2066 (2016).

18 Potapenko, E. S., Biancardi, V. C., Florschutz, R. M., Ryu, P. D. & Stern, J. E. Inhibitory-excitatory synaptic balance is shifted toward increased excitation in magnocellular neurosecretory cells of heart failure rats. Journal of Neurophysiology 106, 1545–1557 (2011).

19 Price, J. F. et al. Arginine Vasopressin Levels Are Elevated and Correlate With Functional Status in Infants and Children With Congestive Heart Failure. Circulation 109, 2550–2553 (2004). Doi: 10.1161/01.CIR.0000129764.84596.EB

20 Patel, T. A., Katsurada, K., Zheng, H. & Patel, K. P. Degradation of nNOS in the PVN of Rats With Heart Failure: Role of CHIP. Hypertension (2025).

21 Roy, R., Wilcox, J., Webb, A. J. & O’Gallagher, K. Dysfunctional and dysregulated nitric oxide synthases in cardiovascular disease: mechanisms and therapeutic potential. International journal of molecular sciences 24, 15200 (2023).

22 Gupta, D. et al. Dietary sodium intake in heart failure. Circulation 126, 479–485 (2012).

23 Mullens, W. et al. Dietary sodium and fluid intake in heart failure. A clinical consensus statement of the Heart Failure Association of the ESC. European Journal of Heart Failure 26, 730–741 (2024).

24 Strazzullo, P., D’Elia, L., Kandala, N.-B. & Cappuccio, F. P. Salt intake, stroke, and cardiovascular disease: meta-analysis of prospective studies. BMJ 339 (2009).

25 Appel, L. J. The effects of dietary factors on blood pressure. Cardiology clinics 35, 197–212 (2017).

26 Ueta, Y. et al. Transgenic expression of enhanced green fluorescent protein enables direct visualization for physiological studies of vasopressin neurons and isolated nerve terminals of the rat. Endocrinology 146, 406–413 (2005). 10.1210/en.2004-0830

27 Althammer, F. et al. Impaired oxytocin signalling in the central amygdala in rats with chronic heart failure. The Journal of Physiology 602, 6259–6280 (2024).

28 Ludwig, M. & Leng, G. Autoinhibition of supraoptic nucleus vasopressin neurons in vivo: a combined retrodialysis/electrophysiological study in rats. European Journal of Neuroscience 9, 2532–2540 (1997).

29 Leng, G. et al. Responses of magnocellular neurons to osmotic stimulation involves coactivation of excitatory and inhibitory input: an experimental and theoretical analysis. Journal of Neuroscience 21, 6967–6977 (2001).

30 Horn, T. F. & Engelmann, M. In vivo microdialysis for nonapeptides in rat brain—a practical guide. Methods 23, 41–53 (2001).

31 Perkinson, M. R. et al. α-Melanocyte-stimulating hormone inhibition of oxytocin neurons switches to excitation in late pregnancy and lactation. Physiological reports 10, e15226 (2022).

32 Chhatbar, P. Y. & Kara, P. Improved blood velocity measurements with a hybrid image filtering and iterative Radon transform algorithm. Frontiers in neuroscience 7, 106 (2013).

33 Schindelin, J., et al. Fiji: an open-source platform for biological-image analysis. Nat Methods 9, 676-682 (2012). 10.1038/nmeth.2019

34 Pitra, S., Zhang, M., Cauley, E. & Stern, J. E. NMDA receptors potentiate activity-dependent dendritic release of neuropeptides from hypothalamic neurons. The Journal of physiology 597, 1735–1756 (2019).

35 Son, S. J. et al. Dendritic peptide release mediates interpopulation crosstalk between neurosecretory and preautonomic networks. Neuron 78, 1036–1049 (2013).

36 Ambach, G. & Palkovits, M. The blood supply of the hypothalamus in the rat. Handbook of the Hypothalamus 1, 267–377 (1979).

37 Kim, J. K. et al. Arginine vasopressin gene expression in chronic cardiac failure in rats. Kidney international 38, 818–822 (1990).

38 Chen, X., Lu, G., Tang, K., Li, Q. & Gao, X. The secretion patterns and roles of cardiac and circulating arginine vasopressin during the development of heart failure. Neuropeptides 51, 63–73 (2015).

39 Belardinelli, L. et al. The A2A adenosine receptor mediates coronary vasodilation. The Journal of pharmacology and experimental therapeutics 284, 1066–1073 (1998).

40 Guerrero, A. A2A adenosine receptor agonists and their potential therapeutic applications. An update. Current medicinal chemistry 25, 3597–3612 (2018).

41 Coney, A. & Marshall, J. M. Role of adenosine and its receptors in the vasodilatation induced in the cerebral cortex of the rat by systemic hypoxia. The Journal of physiology 509, 507–518 (1998).

42 Phillis, J. W. Adenosine and adenine nucleotides as regulators of cerebral blood flow: roles of acidosis, cell swelling, and K ATP channels. Critical Reviews™ in Neurobiology 16 (2004).

43 Ko, K. R., Ngai, A. & Winn, H. R. Role of adenosine in regulation of regional cerebral blood flow in sensory cortex. American Journal of Physiology-Heart and Circulatory Physiology 259, H1703–H1708 (1990).

44 Dirnagl, U., Niwa, K., Lindauer, U. & Villringer, A. Coupling of cerebral blood flow to neuronal activation: role of adenosine and nitric oxide. American Journal of Physiology-Heart and Circulatory Physiology 267, H296–H301 (1994).

45 Matsugi, T., Chen, Q. & Anderson, D. R. Adenosine-induced relaxation of cultured bovine retinal pericytes. Investigative ophthalmology & visual science 38, 2695–2701 (1997).

46 Jackson, E. K. et al. Adenosine production by brain cells. Journal of neurochemistry 141, 676–693 (2017).

47 Marciante, A. B. et al. Microglia regulate motor neuron plasticity via reciprocal fractalkine and adenosine signaling. Nature Communications 15, 10349 (2024).

48 Fu, Z. et al. Microglia modulate the cerebrovascular reactivity through ectonucleotidase CD39. Nature Communications 16, 956 (2025).

49 Császár, E. et al. Microglia modulate blood flow, neurovascular coupling, and hypoperfusion via purinergic actions. Journal of Experimental Medicine 219 (2022).

50 Zhu, C. et al. Expression site of P2RY12 in residential microglial cells in astrocytomas correlates with M1 and M2 marker expression and tumor grade. Acta Neuropathologica Communications 5, 1–12 (2017).

51 Lin, S.-S., Tang, Y., Illes, P. & Verkhratsky, A. The safeguarding microglia: central role for P2Y12 receptors. Frontiers in Pharmacology 11, 627760 (2021).

52 Light, A. R., Wu, Y., Hughen, R. W. & Guthrie, P. B. Purinergic receptors activating rapid intracellular Ca2+ increases in microglia. Neuron glia biology 2, 125–138 (2006).

53 Yu, T. et al. P2Y12 regulates microglia activation and excitatory synaptic transmission in spinal lamina II neurons during neuropathic pain in rodents. Cell death & disease 10, 165 (2019).

54 Swiatkowski, P. et al. Activation of microglial P2Y12 receptor is required for outward potassium currents in response to neuronal injury. Neuroscience 318, 22–33 (2016).

55 Pitra, S., Zhang, M., Cauley, E. & Stern, J. E. NMDA receptors potentiate activity-dependent dendritic release of neuropeptides from hypothalamic neurons. J Physiol 597, 1735–1756 (2019). 10.1113/JP277167

56 Perkinson, M. R. et al. alpha-Melanocyte-stimulating hormone inhibition of oxytocin neurons switches to excitation in late pregnancy and lactation. Physiol Rep 10, e15226 (2022). 10.14814/phy2.15226

57 Zaelzer, C., Gizowski, C., Salmon, C. K., Murai, K. K. & Bourque, C. W. Detection of activity-dependent vasopressin release from neuronal dendrites and axon terminals using sniffer cells. J. Neurophysiol. 120, 1386–1396 (2018).

58 Roy, R. K. et al. Blood flows from the SCN toward the OVLT within a new brain vascular portal pathway. Science advances 10, eadn8350 (2024).

59 Saper, C. B. & Lowell, B. B. The hypothalamus. Current Biology 24, R1111–R1116 (2014).

60 Hartupee, J. & Mann, D. L. Neurohormonal activation in heart failure with reduced ejection fraction. Nature Reviews Cardiology 14, 30–38 (2017).

61 Manolis, A. A., Manolis, T. A. & Manolis, A. S. Neurohumoral activation in heart failure. International Journal of Molecular Sciences 24, 15472 (2023).

62 Cohn, J. N. The management of chronic heart failure. N. Engl. J. Med. 335, 490–498 (1996).

63 Antunes-Rodrigues, J., De Castro, M., Elias, L. L., Valenca, M. M. & McCann, S. M. Neuroendocrine control of body fluid metabolism. Physiological reviews 84, 169–208 (2004).

64 Bourque, C. W. Central mechanisms of osmosensation and systemic osmoregulation. Nature Reviews Neuroscience 9, 519–531 (2008).

65 Ahmad, A. et al. Role of nitric oxide in the cardiovascular and renal systems. International journal of molecular sciences 19, 2605 (2018).

66 Du, W., Stern, J. E. & Filosa, J. A. Neuronal-derived nitric oxide and somatodendritically released vasopressin regulate neurovascular coupling in the rat hypothalamic supraoptic nucleus. J. Neurosci. 35, 5330–5341 (2015).

67 Dixon, A. K., Gubitz, A. K., Sirinathsinghji, D. J., Richardson, P. J. & Freeman, T. C. Tissue distribution of adenosine receptor mRNAs in the rat. Br. J. Pharmacol. 118, 1461–1468 (1996).

68 Ponnoth, D. S. et al. Absence of adenosine-mediated aortic relaxation in A2A adenosine receptor knockout mice. American Journal of Physiology-Heart and Circulatory Physiology 297, H1655–H1660 (2009).

69 Patel, K. P., Zhang, P. L. & Krukoff, T. L. Alterations in brain hexokinase activity associated with heart failure in rats. *American Journal of Physiology-Regulatory*, Integrative and Comparative Physiology 265, R923–R928 (1993).

70 Vahid-Ansari, F. & Leenen, F. H. Pattern of neuronal activation in rats with CHF after myocardial infarction. American Journal of Physiology-Heart and Circulatory Physiology 275, H2140–H2146 (1998).

71 Iovino, M. et al. Vasopressin in heart failure. Endocrine, Metabolic & Immune Disorders-Drug Targets (Formerly Current Drug Targets-Immune, Endocrine & Metabolic Disorders) 18, 458–465 (2018).

72 Van Wylen, D. G., Sciotti, V. M. & Winn, H. R. in Adenosine and adenine nucleotides as regulators of cellular function 191–202 (CRC Press, 2024).

73 Kusano, Y. et al. Role of adenosine A2 receptors in regulation of cerebral blood flow during induced hypotension. J. Cereb. Blood Flow Metab. 30, 808–815 (2010).

74 Soricelli, A. et al. Effect of adenosine on cerebral blood flow as evaluated by single-photon emission computed tomography in normal subjects and in patients with occlusive carotid disease: a comparison with acetazolamide. Stroke 26, 1572–1576 (1995).

75 Shi, Y., Falck, J. R., Harder, D. R. & Koehler, R. C. Interaction of adenosine and cytochrome P450 epoxygenase pathways in neurovascular coupling during whisker stimulation. J. Cereb. Blood Flow Metab. 25, S161–S161 (2005).

76 O’Regan, M. Adenosine and the regulation of cerebral blood flow. Neurol. Res. 27, 175–181 (2005).

77 Ledent, C. et al. Aggressiveness, hypoalgesia and high blood pressure in mice lacking the adenosine A2a receptor. Nature 388, 674–678 (1997).

78 Shi, Y. et al. Interaction of mechanisms involving epoxyeicosatrienoic acids, adenosine receptors, and metabotropic glutamate receptors in neurovascular coupling in rat whisker barrel cortex. J. Cereb. Blood Flow Metab. 28, 111–125 (2008).

79 Fredholm, B. Adenosine, an endogenous distress signal, modulates tissue damage and repair. Cell Death & Differentiation 14, 1315–1323 (2007).

80 Matyash, M., Zabiegalov, O., Wendt, S., Matyash, V. & Kettenmann, H. The adenosine generating enzymes CD39/CD73 control microglial processes ramification in the mouse brain. PloS one 12, e0175012 (2017).

81 Ransohoff, R. M. A polarizing question: do M1 and M2 microglia exist? Nature neuroscience 19, 987–991 (2016).

82 Császár, E. et al. Microglia modulate blood flow, neurovascular coupling, and hypoperfusion via purinergic actions. J. Exp. Med. 219, e20211071 (2022).

83 Ding, Z. et al. Emerging roles of microglia in neuro-vascular unit: Implications of microglia-neurons interactions. Front. Cell. Neurosci. 15, 706025 (2021).

84 Mildner, A., Huang, H., Radke, J., Stenzel, W. & Priller, J. P2Y12 receptor is expressed on human microglia under physiological conditions throughout development and is sensitive to neuroinflammatory diseases. Glia 65, 375–387 (2017).

85 Haynes, S. E. et al. The P2Y12 receptor regulates microglial activation by extracellular nucleotides. Nature neuroscience 9, 1512–1519 (2006).

86 Davalos, D. et al. ATP mediates rapid microglial response to local brain injury in vivo. Nature neuroscience 8, 752–758 (2005).

87 Cserép, C. et al. Microglia monitor and protect neuronal function through specialized somatic purinergic junctions. Science 367, 528–537 (2020).

88 Sipe, G. et al. Microglial P2Y12 is necessary for synaptic plasticity in mouse visual cortex. Nature communications 7, 10905 (2016).

89 Gu, N. et al. Microglial P2Y12 receptors regulate microglial activation and surveillance during neuropathic pain. Brain. Behav. Immun. 55, 82–92 (2016).

90 Althammer, F. et al. Angiotensin II–mediated neuroinflammation in the hippocampus contributes to neuronal deficits and cognitive impairment in heart failure rats. Hypertension 80, 1258–1273 (2023).

91 Badoer, E. Microglia: activation in acute and chronic inflammatory states and in response to cardiovascular dysfunction. The international journal of biochemistry & cell biology 42, 1580–1585 (2010).

92 Kang, Y.-M., Zhang, Z.-H., Xue, B., Weiss, R. M. & Felder, R. B. Inhibition of brain proinflammatory cytokine synthesis reduces hypothalamic excitation in rats with ischemia-induced heart failure. American Journal of Physiology-Heart and Circulatory Physiology (2008).

93 Najjar, F., Ahmad, M., Lagace, D. & Leenen, F. H. Role of myocardial infarction-induced neuroinflammation for depression-like behavior and heart failure in ovariectomized female rats. Neuroscience 415, 201–214 (2019).

94 Rana, I. et al. Microglia activation in the hypothalamic PVN following myocardial infarction. Brain research 1326, 96–104 (2010).

95 Althammer, F. et al. Angiotensin-II drives changes in microglia–vascular interactions in rats with heart failure. Communications Biology 7, 1537 (2024).

96 Akaishi, T. et al. High salt diet intake promotes the induction of experimental autoimmune encephalomyelitis by exacerbating neutrophil infiltration and microglial activation. European Journal of Pharmacology, 177982 (2025).

97 Zhang, T. et al. Excess salt intake promotes M1 microglia polarization via a p38/MAPK/AR-dependent pathway after cerebral ischemia in mice. Int. Immunopharmacol. 81, 106176 (2020).

98 Gilman, T. L., Mitchell, N. C., Daws, L. C. & Toney, G. M. Neuroinflammation contributes to high salt intake-augmented neuronal activation and active coping responses to acute stress. International Journal of Neuropsychopharmacology 22, 137–142 (2019).

99 Pocock, J. M. & Kettenmann, H. Neurotransmitter receptors on microglia. Trends Neurosci. 30, 527–535 (2007).

100 Mairesse, J. et al. Oxytocin receptor agonist reduces perinatal brain damage by targeting microglia. Glia 67, 345–359 (2019).

101 Gonzalez, A. & Hammock, E. A. Oxytocin and microglia in the development of social behaviour. Philosophical Transactions of the Royal Society B 377, 20210059 (2022).

102 Panaro, M. A., Benameur, T. & Porro, C. Hypothalamic neuropeptide brain protection: focus on oxytocin. Journal of Clinical Medicine 9, 1534 (2020).

103 Inoue, T. et al. Oxytocin suppresses inflammatory responses induced by lipopolysaccharide through inhibition of the eIF-2α–ATF4 pathway in mouse microglia. Cells 8, 527 (2019).

104 Pannell, M., Szulzewsky, F., Matyash, V., Wolf, S. A. & Kettenmann, H. The subpopulation of microglia sensitive to neurotransmitters/neurohormones is modulated by stimulation with LPS, interferon-γ, and IL-4. Glia 62, 667–679 (2014).

105 Kisler, K., Nelson, A. R., Montagne, A. & Zlokovic, B. V. Cerebral blood flow regulation and neurovascular dysfunction in Alzheimer disease. Nature Reviews Neuroscience 18, 419–434 (2017).

106 Wolters, F. J. et al. Cerebral perfusion and the risk of dementia: a population-based study. Circulation 136, 719–728 (2017).

