## Supplementary figures and images for "Microglial Purinergic Signaling Underlies Salt-Induced Neurovascular Polarity Reversal in the Hypothalamus During Heart Failure"

### Supplemental Figure 1

#### Supp Figure 1

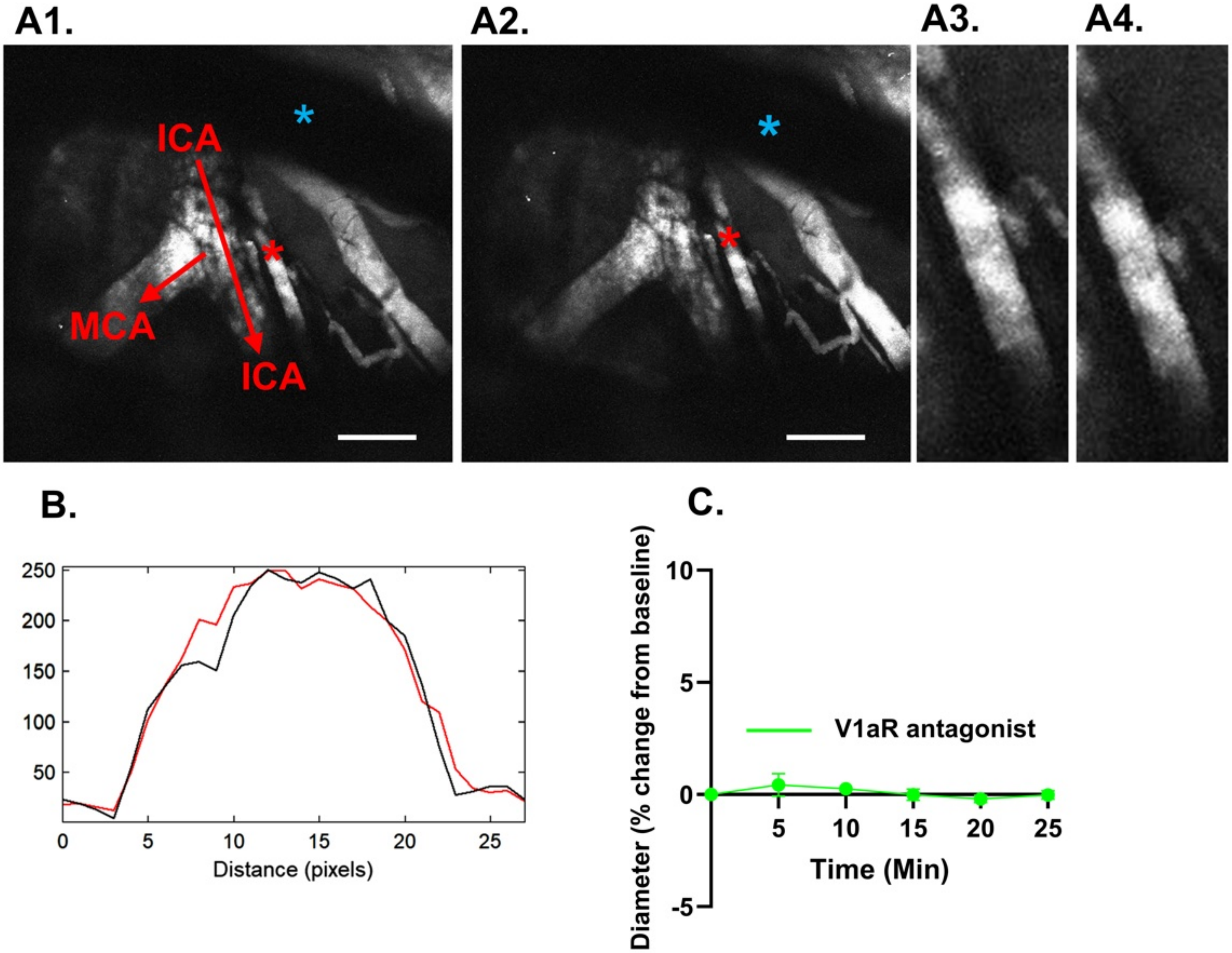
